# Long-Term Potentiation Induces Extrasynaptic Exocytosis of Glun2A-containing NMDA Receptors that is Mainly Controlled by SNAP23

**DOI:** 10.1101/746404

**Authors:** Xiaojun Yu, Wei Li, Tong Wang

## Abstract

NMDA receptors (NMDAR) are key players in the initiation of synaptic plasticity that underlies learning and memory. Long-term potentiation (LTP) of synapses require an increased calcium current via NMDA channels to trigger modifications in postsynaptic density (PSD). It is generally believed that the amount of NMDARs on the postsynaptic surface remains stationary, whereas their subunit composition is dynamically fluctuated during this plasticity process. However, the molecular machinery underlying this subunit-specific regulation remains largely elusive. Here, by detecting the time-lapse changes of surface GluN2A and GluN2B subunit levels using biochemical approaches, surface immunostaining, live-imaging and super-resolution microscopy, we uncovered a transient increase of surface GluN2A-type NMDARs shortly after the induction of chemical long term potentiation (cLTP). These augmented sub-diffraction-limited GluN2A clusters predominantly exist in extrasynaptic domains. We also showed that the spine-enriched SNARE associated protein SNAP-23, and to a minor extent its homologue SNAP-25, control both the basal and regulated surface level of GluN2A receptors. Using a total internal reflection fluorescence microscopy (TIRFM) based live-imaging assay, we resolved and analyzed individual exocytic events of NMDARs in live neurons and found that cLTP raised the frequency of NMDAR exocytosis at extrasynaptic regions, without altering the duration or the package size of these events. Our study thereby provides direct evidence that synaptic plasticity controls the postsynaptic exocytosis machinery, which induces the insertion of more GluN2A receptors into the extrasynaptic area.

**Significance Statement:** Memory formation involves the long-term modification of synapses, which is called synaptic plasticity. In the postsynaptic density (PSD) of excited neurons, this modification process occurs on a minute timescale, initiated by the opening of NMDARs that trigger downstream cascades to fix the potentiation (LTP) at specific synapses for longer timescales. Here, using a novel live-imaging assay we resolved the dynamic delivery of NMDARs to the cell surface, and found that only the insertion frequency, not the duration of individual insertion or number of GluN2A subunits each of these NMDAR vesicles contains, was altered during the synaptic potentiation process. We also identified SNAP-23 as the key molecule mediating this activity dependent NMDAR surface delivery. This study provides a novel mechanism of how NMDARs are regulated in the short window to initiate the long-lasting synaptic modifications.

## Introduction

The N-methyl-D-aspartate (NMDA)-type glutamate receptors (NMDARs) are ionotropic glutamate receptors that allow calcium (Ca^2+^) influx into the postsynaptic compartment and subsequently trigger a variety of intracellular signaling cascades underlying synaptic plasticity, including long-term potentiation (LTP), which has long been postulated as the cellular basis for learning and memory (Tsien et al., 1996; Yashiro and Philpot, 2008). NMDAR channel properties are determined by subunit composition, which governs both their physiological characteristics as well as synaptic distribution and downstream intracellular signalling (Sanz-Clemente et al., 2013). In the forebrain, NMDARs predominantly comprise of two GluN1 subunits and two GluN2A or GluN2B subunits (Kutsuwada et al., 1992). The presence of GluN2A-containing channels (but not GluN2B) is dynamically up-regulated by sensory activation and brain development (Monyer et al., 1994; Barth and Malenka, 2001), and this relative enrichment of GluN2A-containing receptors plays a critical role in synaptic maturation and circuit formation (Yashiro and Philpot, 2008). Recent studies found a significant difference in the surface delivery rate between NMDARs with GluN2A and GluN2B subunits shortly after chemical LTP (cLTP). They established that there is an increase in the surface distribution of GluN2A subunits but not GluN2B (Zhang et al., 2015). The independent regulation of GluN2 subunits underlying synaptic plasticity is further supported by different behavior of their surface nanoclusters, which show distinct patterns in their lateral mobility, distribution and function in live neurons (Bard and Groc, 2011; Ferreira et al., 2017; Kellermayer et al., 2018). However, how neuronal activity shapes distinct distributions of GluN2A- and GluN2B-containing NMDARs between extrasynaptic and synaptic regions is still largely unknown. Therefore, it is critical to investigate activity-regulated machinery underlying surface insertion of NMDARs during synaptic plasticity in order to address the critical question of whether and how is surface addition of these two subunits independently regulated.

All GluN subunits of NMDARs are assembled in the endoplasmic reticulum (ER) then processed and packaged into vesicles throughout the Golgi apparatus. These vesicles carry NMDARs to dendritic spines via long-range intracellular trafficking (Horak et al., 2014). These NMDARs are eventually inserted into the postsynaptic plasma membrane via SNARE (soluble N-ethylmaleimide–sensitive factor attachment receptor) complex-mediated exocytosis (Lledo et al., 1998; Barth and Malenka, 2001; Malinow and Malenka, 2002; Jurado et al., 2013; Horak et al., 2014). SNARE complex is comprised of one copy of vesicle SNARE (*v-*SNARE), such as synaptobrevin/VAMP2, that binds to two counterparts of target SNARE (*t-*SNARE), such as syntaxins and SNAPs, to form a ternary bundle mediating the fusion between vesicles and the plasma membrane (Osborn and Castro, 1977). SNAP-25, a neuron-specific *t-*SNARE, plays a critical role in synaptic vesicle release in presynaptic nerve terminals (Rizo and Sudhof, 2002). SNAP-23 is more ubiquitously-expressed than the SNAP-25 homologue, which mediates vesicle release in non-neuronal cells (Foster et al., 1999; Abonyo et al., 2004; Hepp et al., 2005; Cocucci et al., 2008; Naskar and Puri, 2017). Recently, SNAP-25 was also found to mediate the consecutive insertion of NMDARs in postsynaptic densities (Suh et al., 2010; Jurado et al., 2013; Gu and Huganir, 2016). SNAP-25, but not SNAP-23, is known to be involved in regulated exocytosis of α-amino-3-hydroxy-5-methyl-4-isoxazolepropionic acid receptors (AMPARs) via SNARE-mediated fusion (Jurado et al., 2013), while both SNAP-23 and SNAP-25 have been found to play specific roles in the consecutive trafficking of NMDARs to postsynaptic regions (Lau et al., 2010; Suh et al., 2010; Gu and Huganir, 2016). However, the role of SNAP-23 and SNAP-25 in the activity-regulated exocytosis of GluN2 subunits in postsynaptic neurons is still unknown.

In this study, we addressed these questions by exploring the function of SNAP-23 and SNAP-25 in the activity-dependent insertion of GluN2 subunits. We first identified the activity-dependent surface fluctuations of GluN2A and GluN2B levels by measuring their time-lapse surface intensities using both surface biotinylation and antibody feeding assays in live neurons, and found a transient increase in surface GluN2A-type NMDARs shortly after the induction of cLTP. We next identified that these surface clusters of GluN2A receptors colocalize extensively with SNAP-23 and these augmented GluN2A clusters exist in the extrasynaptic domains of postsynaptic neurons. Then, using botulinum neurotoxin type A (BoNT/A) cleavage as well as their specific shRNAs, we specifically down-regulated the endogenous level of SNAP-23, SNAP-25 or both, and found SNAP-23 is the major regulator underlying both the basal (consecutive) and activity-dependent (regulated) insertion of GluN2A receptors into the plasma membrane following cLTP. This was achieved predominantly by controlling the fusion frequency of vesicles containing GluN2A subunits onto the extrasynaptic sites, which was revealed using a TIRF-based live-imaging method that captures and identifies the properties of individual NMDAR exocytotic events. Our study provides the first direct evidence that synaptic plasticity controls SNAP-23 and hence the postsynaptic exocytosis machinery, which promotes the insertion of GluN2A-containing NMDARs into extrasynaptic regions.

## Material and Methods

### Antibodies, molecular reagents and DNA constructs

Rabbit anti-GluN2A N-terminus polyclonal antibody was obtained from Invitrogen (#480031) and mouse anti-GluN2B N-terminal monoclonal antibodies from NeuroMab (#75-101). Sheep anti-SNAP-23 was obtained from R&D (#AF6306) and mouse anti-SNAP-25 from Synaptic System (#111-011). Mouse anti-Homer1 C-terminal antibody was from NeuroMab (#73-492). Alexa Fluor secondary antibodies were purchased from Life Technologies. DNA constructs encoding BoNT/A-Lc and BoNT/E-Lc were kindly provided by Professor Min Dong (Harvard Medical School, Boston). shRNA sequences against rat SNAP-23 (5’-AATGATGCCAGAGAAGATGAG-3’) as reported (Suzuki and Verma, 2008), and shRNA construct against rat SNAP-25 (5’-GGACGCAGACATGCGTAAT-3’) as reported (Wang et al., 2014) were used. The remaining reagents were obtained from Electron Microscopy Sciences or Sigma Aldrich unless otherwise specified.

### Neuronal cultures and immuno-surface labeling of GluN2 receptors

Hippocampal neurons were cultured from embryonic day 18 (E18) embryos from Sprague Dawley rats of either sex. Animal handling procedures were approved by The University of Queensland Animal Ethics Committee and were conducted in accordance with guidelines set by the Australian National Health and Medical Research Council. Hippocampal neurons were prepared as described previously (Wang et al., 2016) and were plated on either glass coverslips (for confocal microscopy), plastic 12-well plates (for biotinylation) or in glass bottom dishes (Cellvis, #D29-10-1.5-N). For cLTP stimulation, DIV22 rat hippocampal neurons were first incubated in Buffer A (25 mM HEPES, 120mM NaCl, 5mM KCl, 2mM CaCl_2_, 2mM MgCl_2_, 30mM Glucose, pH 7.4, freshly supplemented with 1 μm Strychnine, 0.5 μm TTX) for 1h-2h at 37°C, then treated with prewarmed Buffer B (25 mM HEPES, 120mM NaCl, 5mM KCl, 2mM CaCl_2_, 30mM Glucose, pH 7.4, freshly supplemented with 1 μm Strychnine, 20 μm Bicuculine and 200 μm Glycine) for indicated time. For selective NMDAR antagonist APV treatment, neurons were preincubated in Buffer A supplemented with APV (50 μM) for 30 min and then stimulated with Buffer B in the presence of APV (50 μM) for the indicated amount of time. For GluN2A and GluN2B surface immunofluorescence staining, antibodies against the extracellular N-terminus of both subunits were diluted at 1:200 dilution using cold buffer A. Neurons blocked with antibody solution were kept on ice for 30 min, then fixed in 4% paraformaldehyde and 4% sucrose in PBS for 20 min at room temperature, and incubated with Alexa-647 conjugated secondary antibodies for 1h in room temperature. Neurons were next washed and permeabilized with 0.1% Triton X-100 in PBS, incubated with other primary antibodies, then with fluorescence-conjugated secondary antibodies, and mounted in liquid mounting medium (Vectashield, H-1200) for imaging using either confocal or structured illumination microscope (SIM). Image analysis was carried out using Zen (Carl Zeiss), Image J (NIH) or Imaris (Bitplane) software, with subsequent quantification performed using Prism (Graphpad). Colocalization Coefficient *R_coloc_* (Zinchuk et al., 2007), is described by the equation (1), using the ImageJ plugin JACoP (Bolte and Cordelieres, 2006).

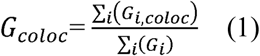

where *G_i_* refers to the intensity values of the GFP channels of pixel *i*. *G_i,coloc_* is the pixel Colocalized with Homer1c-DsRed channel. The ratio of extrasynaptic GluN2A receptors was calculated using the following equation:

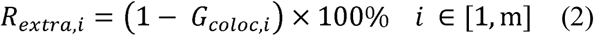

where *i* refers to the *i^th^* analysed image out of the total of *m* images.

### Biotinylation labeling and western detection of GluN2 receptors

DIV22 rat cortical neurons cultured in 12-well plates were treated with cLTP as described above, then incubated with 0.5 mg/ml nonpermeable biotin (NHS-biotin, Sigma-Aldrich) in cold PBS for 30 min under gentle agitation. After incubation, cells were washed with TBS three times, total wash time 30 min. Cell samples were then centrifuged at 14,000*g* for 15 min, and the supernatant collected and added to streptavidin-agarose beads (Thermo Fisher Scienctific) and rotated overnight at 4°C. The beads were washed twice with wash buffer containing 150 mM NaCl, 50 mM HEPES, 0.1% Triton X-100, then boiled at 70°C for 5 min using sample loading buffer (250 mM Tris–HCl (pH 6.8), 8% (w/v) sodium dodecyl sulfate (SDS), 0.2% (w/v) bromophenol blue, 40% (v/v) glycerol, 20% (v/v) β-mercaptoethanol), and the supernatant was collected for western blot detection of the surface GluN2 subunit levels.

### Bleaching assay for GluN2A receptor exocytosis

For all live-imaging experiments, DIV21-35 rat hippocampal neurons were used. Time-lapse imaging of super-ecliptic pHluorin–tagged GluN2A (SEP-GluN2A) exocytosis was performed at 37°C in live-imaging buffer (25 mM HEPES, 120mM NaCl, 5mM KCl, 2mM CaCl_2_, 2mM MgCl_2_, 30mM Glucose, 0.5 µM Strychnine and 0.5 µM TTX, pH 7.4). We employed Elyra SIM/PALM/STORM Microscope equipped with Andor 897 EMCCD camera suitable for single-molecule imaging to visualize insertion of individual vesicles containing NMDARs and detect single SEP-GluN2A subunits. The microscope was used in total internal reflection fluorescence (TIRF) mode. Before imaging of exocytic fusion events, dual-color image of SEP-GluN2A and Homer 1c-DsRed was first acquired in transfected neurons by sequentially imaging SEP and DsRed channels using 488 nm and 561 nm laser excitations, respectively. Then, SEP-GluN2A was bleached using 488 nm laser at 20 mW laser intensity for 1 min. In the bleached regions, time-lapse imaging of SEP-GluN2A exocytosis was captured for 5 min, using 488 nm laser excitation at 1 mW laser intensity. Measurement of the fusion events induced by cLTP in cultured neurons was performed after bleaching. The number, duration and package size of these exocytotic events was analyzed using methods established in this study, as illustrated in Fig. 5C and described in the next section.

### Analysis of SEP-GluN2A exocytosis

Exocytosis frequency of SEP-GluN2A was analyzed using the TrackMate plugin (Version 3.4.2) of Image J (NIH). To optimize the signal to noise ratio, time-lapse images were first processed in Image J using the following steps. First, regions of interest (ROI) of SEP-GluN2A transfected neurons were manually traced from the images acquired before bleaching. Then in the post-bleaching movies, signal to noise ratio of time-lapse images was further optimized using the filter based on the standard deviation of 4 consecutive images, using the following equation:

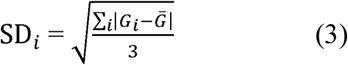

where Ḡ refers to the averaged intensity values of the green channel, *i* refers to the number of pixels. The resultant movies was then subjected to Gaussian blur filter (δ=2 pixel), with background noise removed using the ‘subtract background’ function (Rolling ball radius = 10 pixel). Finally, the movies were traced with TrackMate, and only fusion events with track duration over 1 s quantified.

### Structured illumination microscopy and Imaris Analysis

Imaging of fixed samples was performed using the Elyra SIM/PALM/STORM Microscope (Carl Zeiss) equipped with a 100x objective (α Plan-Apochromat 100×/1.46 oil-immersion DIC M27) and a PCO sCMOS camera (PCO Scientific). Images were obtained by acquiring z-stacks of 10-16 slices with spacing of 0.101 μm and exposure time of 100 ms, SIM grating size of 42-51 μm, and using five rotations. 3D structured illumination images were then aligned and processed using the Zen software. The resultant 3D-SIM images were then analysed using surface function of Imaris software (Bitplane).

### Counting of the number of GluN2A subunits in exocytic vesicles

For single-step photobleaching analysis, time-lapse images of discrete exocytic fusion events were analysed using a custom written Python routine as previously described (Durisic et al., 2014). Briefly, a square region of 3×3 pixels around each fusion event was used to extract an intensity-time profile. The background was calculated locally from a 1 pixel thick frame around the region and subtracted from the signal. Spots with signal to noise ratios less than 2, those that were fluorescent in the first frame of the image sequence and spots that lasted less than five frames were excluded from the analysis as we couldn’t reliably count the number of steps from their intensity-time traces (Li et al., 2016). The steps in the traces were counted manually. In less than 1% of traces, two fluorescent proteins photobleached simultaneously (missed events) and we corrected for those (Durisic et al., 2012).

### Experimental design and statistical analysis

We used GraphPad Prism 7 (GraphPad Inc.) for statistical analysis. Results are reported as mean ± s.e.m. For group comparisons, two-tailed nonparametric *t*-tests or paired *t*-tests were executed. P values < 0.05 indicated statistical significance. No statistical methods were used to predetermine sample sizes. There was no formal randomization. Data collection and analysis were performed by different operators, who were blind to the conditions of the experiments.

## Results

### cLTP induces a transient up-regulation of surface GluN2A-NMDA receptors in cultured mature hippocampal neurons

To observe the effect of synaptic plasticity on the surface levels of NMDA receptors, we potentiated the postsynaptic neuron using a Glycine-induced cLTP protocol. This protocol creates NMDA-dependent LTP (Lu et al., 2001). As shown in the schematic diagram (Fig1. A), mature rat hippocampal neurons (DIV21-28) were first pretreated with inhibitory Buffer A for 1 hour to suppress the intrinsic synaptic activity, then briefly stimulated with 10 min treatment of Buffer B, a Mg^2+^-free buffer that contains 200 nM Glycine to activate the synaptic NMDARs which are receiving spontaneous release of glutamate (Fortin et al., 2010). The efficacy of this protocol was further validated by imaging the subcellular Ca^2+^ influx in neurons expressing the genetically-encoded Ca^2+^ indicator, GCaMP6-fast variant, which has previously been used to label physiological levels of fast Ca^2+^ fluctuations (Chen et al., 2013; Lock et al., 2015). Neurons treated with Buffer A showed a low level of fluctuation in Ca^2+^ within postsynaptic spines, whereas the addition of Buffer B significantly boosted the intensity of this signal (Fig. 1B-D; sMovie 1). This surge in Ca^2+^ signal was totally abolished by the addition of specific NMDA channel antagonist APV (Fig. 1E). These results suggest the cLTP protocol outlined in Figure 1A was able to induce cLTP in cultured rat hippocampal neurons.

**Figure 1.**
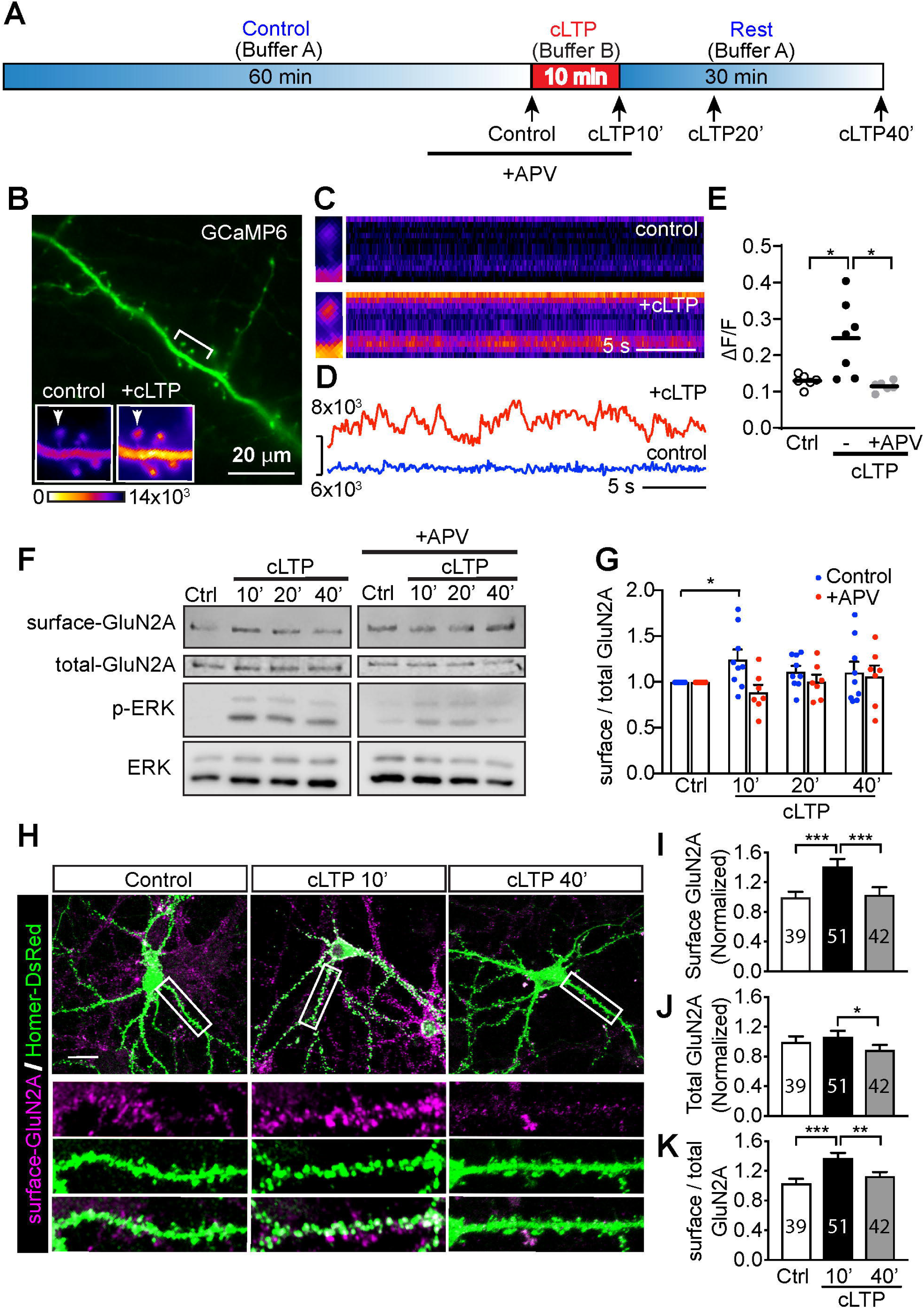
cLTP induces a transient increase in surface GluN2A NMDA receptors in hippocampal neurons. **(A)** Schematic diagram of the cLTP model used in this study. Mature hippocampal neurons (DIV>21) were pre-incubated in Buffer A for 60 min and then flushed with 200 nM of Glycine (Buffer B) for 10 min to induce glycine-dependent cLTP. They were then returned to resting condition in Buffer A for 10 to 30 min. 50 μM APV was added to the bathing medium as indicated. Surface levels of GluN2 receptors were detected at indicated time points. **(B)** Calcium fluctuations of dendritic spine regions visualized in GCaMP6-expressing hippocampal neurons, which were treated with the above cLTP protocol. Fluorescence intensity of bracketed area is amplified in lower left boxes. Bar=20 μm. Kymograph **(C)** and intensity profile **(D)** of the Ca^2+^ intensity in individual dendritic spines before (control) and during the 10 min of cLTP treatment (+cLTP). **(E)** Quantification of Ca^2+^ fluctuations induced by the cLTP protocol. ⊗F/F of the GCaMP6-expressing neurons was used to show the Ca^2+^ intensity change before (control), and during the cLTP treatment without (-) or with (+APV) the existence of 50 μM APV. Results are shown in scatter plot with mean±SEM, \**p*<0.05. n=6 (control), 7 (cLTP), 6 (+APV) cells from two independent cultures, two tailed student’s *t*-test. **(F)** Representative western blots showing the levels of biotin labeled surface GluN2A receptors in cultured rat cortical neurons treated as indicated. **(G)** Quantification of results in **(F)**. Results are shown in scatter plot with mean±SEM, \**p*<0.05. n=9 (ctrl), 7 (cLTP) repeats from at least seven independent preparations, two tailed student’s *t*-test. **(H)** Confocal images showing surface immunostaining for GluN2A receptors in cultured rat hippocampal neurons treated as indicated. Bar=20 μm. **(I-J)** Quantification of immunostaining of surface GluN2A receptor levels **(I)**, total GluN2A receptor levels, which is the GluN2A immunostaining signal after cell membrane permeabilization. **(J)** and the ratio of surface to total GluN2A receptor levels **(K)** before or after cLTP treatment for indicated time. Results are shown in mean±SEM, \**p*<0.05, \*\**p*<0.01, \*\*\**p*<0.001. Results were from n=39 (ctrl), 51 (cLTP), 42 (+APV) cells of three independent preparations, two tailed student’s *t*-test.

We next used the cLTP protocol to investigate the dynamic change in surface NMDARs. Surface levels of both GluN2A and GluN2B subunits were examined at four different time points before or after cLTP stimulation, designated as control (0’), 10’, 20’ and 40’ in Fig. 1A. The levels of surface NMDA receptors were examined using biotinylation labelling followed by western blot detection with specific antibodies against GluN2A or GluN2B, respectively (Tan et al., 2017). The surface level of GluN2A subunits was increased following 10 min of cLTP (Fig. 1F-G, 10’) but was reduced to basal level 30 min post cLTP (Fig. 1F-G, 40’). This transient up-regulation of GluN2A receptors was abolished by APV treatment (Fig. 1F-G, +APV). In contrast, the surface level of GluN2B was not significantly up-regulated following the 10 min cLTP induction (sFig 1A-B). These results show that cLTP differentially controls the synaptic GluN2A and GluN2B subunits by selectively increasing the GluN2A-containing NMDARs insertion following cLTP.

A series of new studies has revealed that NMDARs cycle rapidly into and out of synapses, through either lateral diffusion or new vesicle insertion (Newpher and Ehlers, 2008; Bard and Groc, 2011). To further investigate the dynamic changes in surface levels of GluN2A receptors during LTP, we used an antibody against the N-terminal extracellular domain of GluN2A receptors to examine their surface distribution in live neurons at various time points following cLTP induction. We found that postsynaptic GluN2A receptors colocalized with postsynaptic density (PSD) scaffold protein Homer-1c (Fig. 1H; quantified in Fig. 1I) (Hayashi et al., 2009). This was consistent with the trend shown in Figure 1F, where the surface staining intensity of postsynaptic GluN2A receptors was significantly increased following 10 min cLTP induction, and then reduced to basal level at 40 min. As in our surface biotinylation experiments, total GluN2A level was not significantly increased after 10 min cLTP (Fig. 1J). Similarly, the normalized surface-to-total GluN2A level showed a transient increase at 10 min, which returned to basal level at 40 min (Fig. 1K). This cLTP-induced transient surface expression of GluN2A was also dependent on the opening of NMDA channels, as inhibition with APV was able to totally abolish the surface SEP-GluN2A containing NMDAR increase induced by 10 min cLTP (Fig. 2A, B). The similar cLTP-depedent increase of extrasynaptic surface GluN2A receptors was also observed using antibodies against endogenous GluN2A and Homer-1c (sFig. 2A, B). These results suggest that a local mechanism mediating receptor surface expression was activated by the cLTP treatment.

**Figure 2.**
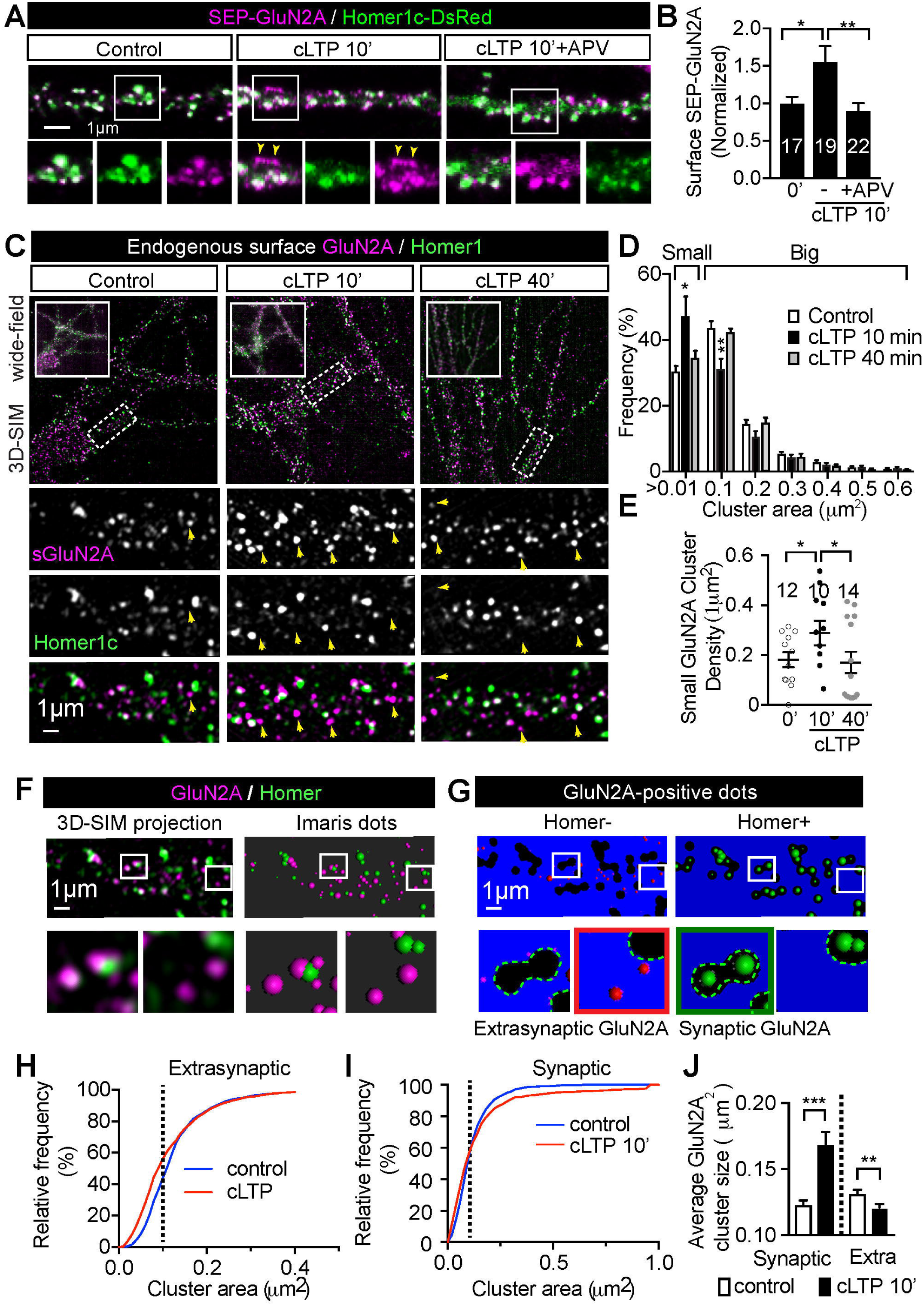
Extrasynaptic GluN2A-NMDARs are up-regulated shortly following the cLTP induction. **(A)** Representative confocal images of surface GluN2A immunostaining in cultured rat hippocampal neurons co-transfected with SEP-GluN2A and Homer1c-DsRed, which were treated as indicated. Boxed regions were amplified in the bottom rows. Extrasynaptic NMDARs were indicated with arrowheads. Bar=1 μm. **(B)** Quantification of immunostaining for surface GluN2A receptor levels. Data represent mean±SEM, n=17 (0’), 19 (-) and 22 (+APV) neurons. \**p*<0.05, \*\**p*<0.01, two-tailed student’s *t*-test, data were derived from three independent cultures. **(C)** Representative 3D-SIM images of endogenous surface GluN2A and Homer immunostaining in cultured rat hippocampal neurons treated as indicated. Wide-field images are shown in the upper left insets. Boxed regions of interest (ROIs) are shown in lower panels. Extrasynaptic NMDARs were indicated with arrowheads. Bar=1 μm. **(D)** Cluster size of Surface GluN2A receptor were determined using the ‘surface’ function of Imaris. The frequency distribution of cluster size were shown in control, cLTP 10 min and cLTP 40 min neurons. Clusters with a diameter of 0.01 to 0.1 μm^2^ were characterized as ‘small’ clusters, whereas all others were characterized as ‘big’ clusters. Data represent mean±SEM, n=6 (control), 5 (10 min) and 7 (40 min) cells were analyzed. \**p*<0.05, \*\**p*<0.01, two-tailed student’s *t*-test, data were derived from three independent cultures. **(E)** Quantification of the average density of small GluN2A clusters. Data represent mean±SEM, n=12 (0’), 10 (cLTP 10’) and 14 (cLTP 40’) neurons. \**p*<0.05, two-tailed student’s *t*-test, data were derived from three independent cultures. **(F)** The dual-colour SIM images were analysed using Imaris software, and the representative Imaris-rendered surfaces were shown. GluN2A and Homer were extracted as separate surfaces to detect the overlap between surface of GluN2A receptors and that of the postsynaptic marker Homer. Boxed regions are amplified in lower panels. Bar=1μm. **(G)** Representative spatial separation efficiency of the Imaris surface method in extracting the synaptic and extrasynaptic GluN2A receptors. Non-Homer1c overlapping (Homer-) GluN2A receptors were characterized as extrasynaptic, and Homer1c overlapping (Homer+) GluN2A receptors were characterized as synaptic. Boxed regions are amplified in the bottom panels with extrasynaptic or synaptic GluN2A receptors marked using red and green surfaces, respectively. **(H-I)** Cumulative frequency distribution of the size of extrasynaptic **(H)** and synaptic **(I)** GluN2A receptors surfaces in cultured hippocampal neurons before (control) or after 10 min of cLTP treatment (cLTP). **(J)** Average cluster sizes of the synaptic and extrasynaptic surface GluN2A receptors, in cultured hippocampal neurons before (control) or after 10 min of cLTP treatment (cLTP). Data represent mean±SEM, n=606, 800, 981 and 1323 clusters from at least five neurons were compared. \*\**p*<0.01, \*\*\**p*<0.001, two-tailed student’s *t*-test, data were derived from three independent cultures.

### cLTP raises the ratio of small GluN2A receptor clusters in the extrasynaptic area

By comparing the distribution of surface GluN2A receptors before and after 10 min cLTP, we also noticed that many of these newly increased GluN2A receptors were in the extrasynaptic regions, and not overlapping with Homer-1c (Fig. 2A, arrows). It has been shown that both the mobility and clustering of surface GluN2A receptors are affected by cLTP, and that clustering of GluN2A receptors in micro- and/or nano-domains of PSD is critical for the function of synaptic plasticity (Bard and Groc, 2011; Ferreira et al., 2017; Kellermayer et al., 2018). We therefore used sub-diffraction structured illumination microscopy (SIM) to characterize the relative distribution of surface GluN2A receptors with respect to Homer-1c, which marks the synapses in postsynaptic neurons, before and after 10 min of cLTP treatment, as shown in Fig. 2C. To describe the distribution of surface GluN2A micro- and nano-clusters, we first compared the distribution of GluN2A cluster size in 50 μm 50 μm ROIs. We found that the ratio of GluN2A receptor clusters with smaller size (< 0.1 μm^2^; Fig. 3D, ‘small’) was significantly increased after 10 min of cLTP treatment (Fig. 2D, ‘40 min’) and reduced to basal level at 30 min after stimulation (Fig. 2D, ‘40 min’). Consistently, the absolute number of small GluN2A clusters was also transiently increased after 10 min of cLTP treatment (Fig. 2E).

**Figure 3.**
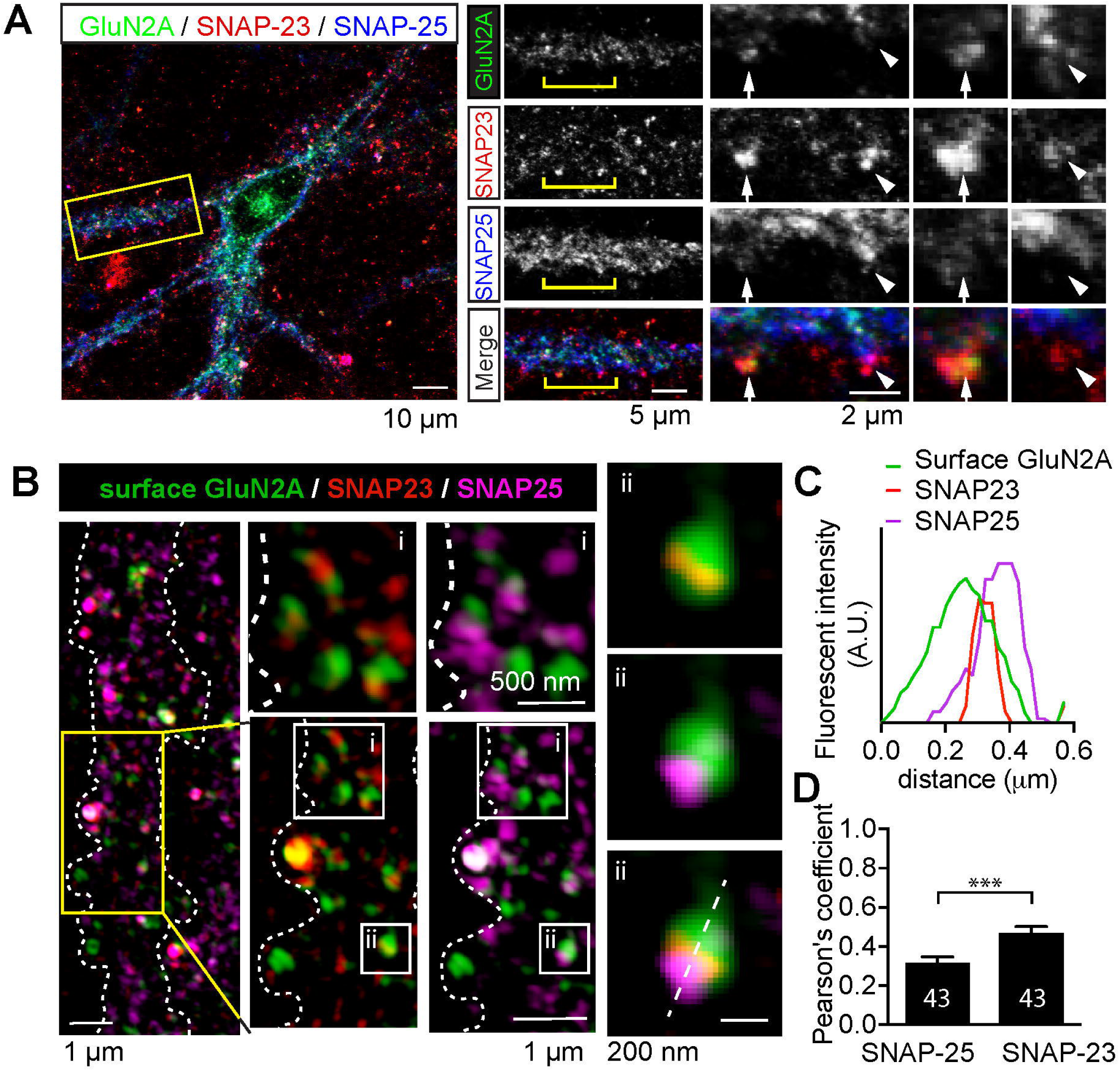
Surface GluN2A clusters are more closely correlated with the postsynaptic SNARE component SNAP-23 than SNAP-25. **(A)** DIV21 rat hippocampal neurons were fixed and stained with specific antibodies against endogenous GluN2A, SNAP-23 and SNAP-25. Confocal microscopy was used to detect the distribution of signals from three different fluorescent channels. To the right boxed region is amplified with channels separated. Bracketed regions are further amplified. Spine annotated by arrows were positive for GluN2A, both SNAP-23 and SNAP-25, whereas spines annotated by arrowheads shows the overlapping between GluN2A and only SNAP-23. Scale bar as indicated on the panels. **(B)** DIV21 rat hippocampal neurons were live-stained for surface GluN2A expression, followed by fixation and staining with specific antibodies against endogenous SNAP-23 and SNAP-25. 3D-SIM was used to visualize the distribution of fluorescent staining signals at dendritic spines. ROIs were amplified in top and right panels, with 3-color line-profile performed across the single synapse shown in right panel. Scale bar as indicated. **(C)** Line profiles showing the colocalization of GluN2A (green), SNAP-23 (red) and SNAP-25 (magenta). **(D)** Colocalization level between GluN2A and SNAP25 or SNAP23 were quantified with Pearson’s coefficient. Data represent mean ± SEM, n=43 ROIs with SNAP-25 and SNAP-23, respectively. \*\*\**p*<0.001, two-tailed student’s *t*-test, data were measured from two independent cultures.

To further define whether cLTP treatment differently impacts endogenous GluN2A receptor clusters in synaptic and extrasynaptic areas, we extracted and analyzed clusters of synaptic and extrasynaptic GluN2A receptors, respectively. To achieve this, we marked the GluN2A clusters that overlapped the PSD marker Homer-1c as synaptic, whereas the non-Homer-1c-overlapping clusters were defined as the extrasynaptic GluN2A receptors (Fig. 2F-G). The cumulative frequency plot of both extrasynaptic and synaptic GluN2A receptors were compared in control or 10 min cLTP treated neurons. As expected, a significantly increased ratio of extrasynaptic GluN2A receptors was found in small-sized fractions, as reflected by the cumulative frequency distribution (Fig. 2H). This is consistent with the increased frequency distribution of small-sized GluN2A clusters in the total population. Moreover, we also detected an increase in the number of very large (∼1 μm^2^) synaptic GluN2A clusters (Fig. 2I). This is likely to be caused by the increased spine number / size which signifies the structural plasticity caused by the cLTP treatment (Hruska et al., 2018). In agreement with this, the cluster size of the synaptic GluN2A clusters is significantly increased, whereas the size of the extrasynaptic GluN2A clusters exhibit is reduced (Fig. 2J). These results suggest that 10 min of cLTP treatment causes an increase in the amount of GluN2A receptors in the form of small clusters (< 0.1 μm^2^ in area) in the extrasynaptic regions of postsynaptic neurons, in addition there is an increase in the frequency of large clusters in the synaptic regions.

### The cLTP-induced transient increase in GluN2A-NMDAR surface levels is dependent on activity-dependent exocytosis machinery

Insertion of NMDARs to the postsynaptic membrane are mediated by SNARE complexes (Suh et al., 2010; Jurado et al., 2013), in which SNAP23 and SNAP-25 are core components (Osborn and Castro, 1977; Rizo and Sudhof, 2002). Both SNAP-23 and SNAP-25 have been implicated in LTP triggered AMPA receptor surface expression (Jurado et al., 2013). SNAP-23, but not SNAP-25, is required for maintaining basal levels of GluN2A NMDA receptor surface expression in the mouse brain (Suh et al., 2010). We therefore sought to investigate the role of SNAP-23 and SNAP-25 in LTP induced GluN2A receptor surface expression. We analyzed the relative location between GluN2A receptors and SNAP-23 or SNAP-25 in the postsynaptic compartment of neurons and found that both SNAP-25 and SNAP-23 are distributed in the spines, with SNAP-25 more broadly distributed both pre- and post-synaptically, suggesting its function in both presynaptic vesicle recycling (Osborn and Castro, 1977; Rizo and Sudhof, 2002) and postsynaptic glutamate receptor exocytosis (Suh et al., 2010; Jurado et al., 2013). In contrast, SNAP-23 localized specifically to the tips of the postsynaptic spines, which are also enriched with GluN2A receptors (Fig. 3A). These observations are similar to previous studies (Suh et al., 2010; Jurado et al., 2013), suggesting a possible role for SNAP-23 in regulating postsynaptic exocytosis. To further explore the distribution of endogenous SNAP-23 and SNAP-25 in postsynaptic densities, we used subdiffractional SIM to dissect the nano-scale localization of SNAP-23, SNAP-25 and surface clustering of GluN2A receptors simultaneously. As shown in Fig. 3B, we found SNAP-23 (red) more closely correlated to clusters of surface GluN-2A receptors then SNAP-25 (magenta). A larger fraction of SNAP-25 did not overlap with GluN2A receptor patch (Fig. 3C). The overall higher colocalization between GluN2A receptor and SNAP-23 was confirmed using the Pearson’s colocalization coefficient (Fig. 3D). These results suggest that SNAP-23, which has higher postsynaptic specificity, is more likely to regulate the surface expression of GluN2A receptors in postsynaptic spines than the more broadly distributed SNAP-25.

We next explored the role of SNAP-23 and SNAP-25 using botulinum neurotoxins, which cleave and inactivate components of SNARE complex at different sites (Schiavo et al., 2000; Meunier et al., 2003). We used the catalytic light chain of botulinum neurotoxin type A (BoNT/A-Lc), which inactivates both SNAP-23 and SNAP-25 by cleaving their last 9 amino acid residues, as well as the light chain of botulinum neurotoxin type E (BoNT/E-Lc), which at nanomolar concentration only cleaves SNAP-25 (but not SNAP-23) at the last 26 C-terminal residues (Vaidyanathan et al., 1999). We first examined the cleavage of BoNT/A-Lc on both SNAP-25 and SNAP-23 by co-expressing GFP-tagged rat SNAP-23 or SNAP-25 with either BoNT/A-Lc or BoNT/E-Lc in HEK293 cells respectively, and found that both SNAP-25 and SNAP-23 are cleaved by BoNT/A-Lc as reported (Vaidyanathan et al., 1999) (Fig. 4A). We also showed the cleavage of SNAP-25 but not SNAP-23 by BoNT/E-Lc, as previously described (Peng et al., 2013) (Fig. 4A). These results confirmed the effectiveness of both BoNT/A-Lc and BoNT/E-Lc as a tool to inactivate endogenous SNAP-25 and SNAP-23. We then used this tool to manipulate the level of endogenous SNAP-23 and SNAP-25 to observe the impact on cLTP-dependent surface expression of GluN2A receptors in cultured hippocampal neurons. Levels of endogenous GluN2A receptors before and after 10 min of cLTP was examined in neurons co-transfected with BoNT/A-Lc or BoNT/E-Lc and Homer1c-DsRed. BoNT/A-Lc significantly decreased the surface level of GluN2A at resting conditions, and also abolished the cLTP-dependent increase in surface GluN2A expression (Fig. 4B, C). However, the over-expression of BoNT/E-Lc, which truncates a larger fragment from SNAP-25 (Vaidyanathan et al., 1999; Meunier et al., 2003; Peng et al., 2013), caused the degeneration of transfected neurons as previously reported (sFig. 3A) (Peng et al., 2013). We therefore used BoNT/A-Lc as the tool to suppress the activity of endogenous SNAP-25 and SNAP-23 in mature neurons. To explore whether the activity of SNAP-23 or SNAP-25 is relevant to the surface expression of GluN2A receptors, we also used shRNA to knock down (KD) endogenous SNAP-23 or SNAP-25 to investigate their impact on the cLTP-dependent surface expression of GluN2A receptors. The KD efficiency of specific shRNAs against endogenous SNAP-23 or SNAP-25 was tested first as shown in Fig. 4D. We found that KD of either SNAP-25 or SNAP-23 significantly decreased the basal level of GluN2A at resting conditions and abolished the cLTP-dependent increase in surface GluN2A expression (Fig. 4E, F). In contrast, in shSNAP-25 transfected neurons, the surface expression of GluN2A still showed an up-regulated trend in response to cLTP treatment. Its basal level was reduced to 56.54±6.09%, compared to 35.18±3.01% in the shSNAP-23 group (Fig. 4F). This is indicating that in SNAP-25 down-regulated neurons, both basal- and cLTP-induced GluN2A receptor exocytosis was partially preserved. These data suggest that SNAP-23 and SNAP-25 are both required for constitutive GluN2A receptor delivery to the plasma membrane and further support the notion that the cLTP-induced increase in GluN2A receptor level is more likely to be dependent on SNAP-23 mediated exocytosis then on exocytosis mediated by SNAP-25.

**Figure 4.**
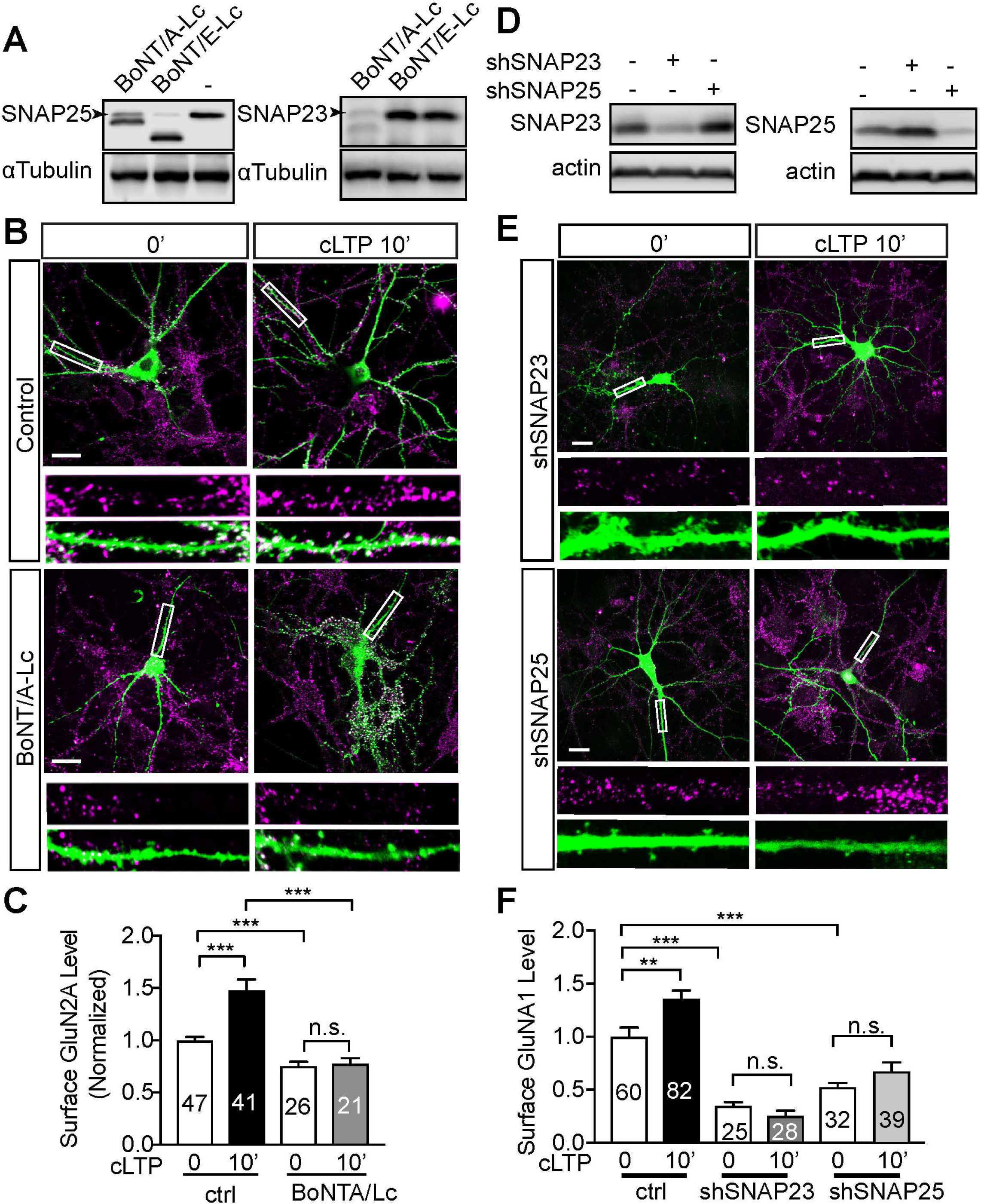
Activity-dependent surface increase in GluN2A receptors is dependent on SNAP-23 and SNAP-25. **(A)** PC12 cells were transfected with plasmids expressing the catalytic light-chains (Lc) of BoNT/A or BoNT/E, respectively. Cleavage of SNAP-23 and SNAP-25 was detected using their specific antibodies, full-length of SNAP-25 and SNAP-23 are indicated with arrows. **(B)** Representative images showing the surface level of endogenous GluN2A following 10 min cLTP stimulation in DIV21-28 rat hippocampal neurons, which were co-transfected with Homer-DsRed and BoNT/A-Lc. Boxed regions showing the level of surface GluN2A subunits were amplified in the bottom panels. Scale bar=20 μm. **(C)** Quantification of the surface level of endogenous GluN2A in BoNT/A-Lc transfected groups. **(D)** PC12 cells were transfected with shSNAP-23 or shSNAP-25 plasmids to verify knock down efficiencies. Endogenous SNAP-23 and SNAP-25 was detected using their specific antibodies. **(E)** Representative images showing the surface level of endogenous GluN2A following 10 min cLTP stimulation in DIV21-28 rat hippocampal neurons co-transfected with Homer-DsRed and shSNAP-23, shSNAP-25 plasmids. Boxed regions showing the level of surface GluN2A subunits were amplified in the bottom panels. Scale bar=20 μm. **(F)** Quantification of the surface level of endogenous GluN2A in shSNAP-23, shSNAP-25 transfected groups. Results are shown in mean±SEM, \*\*\**p*<0.001, n.s. no significant difference. For E, n=47, 41, 26 and 21 cells from three independent preparations, two tailed student’s *t*-test. For F, n=60, 82, 25, 28, 32 and 39 cells from three independent preparations, two tailed student’s *t*-test.

### Exocytosis of GluN2A receptors to the extrasynaptic regions is specifically increased by cLTP

To analyze individual exocytosis of GluN2A-containing NMDARs in postsynaptic neurons, we fused superecliptic pHluorin to the extracellular region of GluN2A (SEP-GluN2A) (Kopec et al., 2006) (Fig. 5A). Using this construct, we were able to detected a >20 fold increase in fluorescence intensity when the acidified intracellular vesicles (pH5.5) fused with the plasma membrane and exposed their SEP units to the neutral extracellular medium (pH>7.0) (Fig. 5B). We further combined this with TIRF microscopy, which only detects GluN2A receptor exocytosis near the plasma membrane (∼100 nm), with a time resolution of ∼10 Hz (Lin et al., 2009; Gu and Huganir, 2016; Joensuu et al., 2017). To precisely examine whether the frequency, duration or location of the regulated GluN2A receptor exocytosis is regulated by cLTP, we employed an additional bleaching step in TIRF mode to remove the pre-existing SEP-GluN2A fluorescence on the plasma membrane. This bleaching step was carried on immediately before the time-lapse acquisition, which was accompanied with the 5 to 10 min of cLTP perfusion (Fig. 5C-D, step ⍰ - ⍰; sFig. 4A). After bleaching, individual exocytic fusion events of vesicles containing SEP-GluN2A receptors could be easily detected near the Homer1c-DsRed-positive postsynaptic compartments (Fig. 5C-D, step ⍰; sFig. 4A; sMovie 2). The projection of time-lapse images of these events on the Homer1c-DsRed channel demonstrated the extrasynaptic location of vesicle fusion in postsynaptic neurons (Fig. 5E), which is similar to previous findings (Newpher and Ehlers, 2008). Next, the time-lapse images of the newly inserted SEP-GluN2A-containing vesicles were automatically traced and both the duration and frequency of these identified trajectories were analyzed (Fig. 5F; see also sFig. 4B-D). As duration of the SEP-GluN2A events reflects the pertinent properties of the underlying exocytotic machinery (Gu and Huganir, 2016), we first compared the average duration of SEP-GluN2A in both control and cLTP conditions and found that they are both around two seconds (Fig. 5G; control: 1.98 ±0.169 s; cLTP: 1.96 ±0.093 s; see also sFig. 3E), with no significant difference between the two treatments (Fig. 5G). This dynamic property of NMDA receptor exocytosis is consistent with previous studies (Gu and Huganir, 2016). However, in contrast to their unchanged duration, the fusion frequency of vesicles contains SEP-GluN2A receptors was significantly increased by the cLTP treatment (Fig. 6A-B). Moreover, the cLTP-induced frequency increase of the SEP-GluN2A insertion was totally blocked by the specific NMDA channel antagonist APV (Fig. 6B). Importantly, we also found that co-expression of BoNT/A-Lc completely abolished cLTP-induced increase of SEP-GluN2A insertion (Fig. 6B), indicating this increased GluN2A insertion was indeed caused by the cLTP-dependent regulation of postsynaptic exocytotic machinery. These results suggest that during the early stage of cLTP, an NMDAR-mediated response up-regulates the frequency of SEP-GluN2A exocytotic insertion into the postsynaptic compartments, without affecting the dynamic properties pertinent to these events.

**Figure 5.**
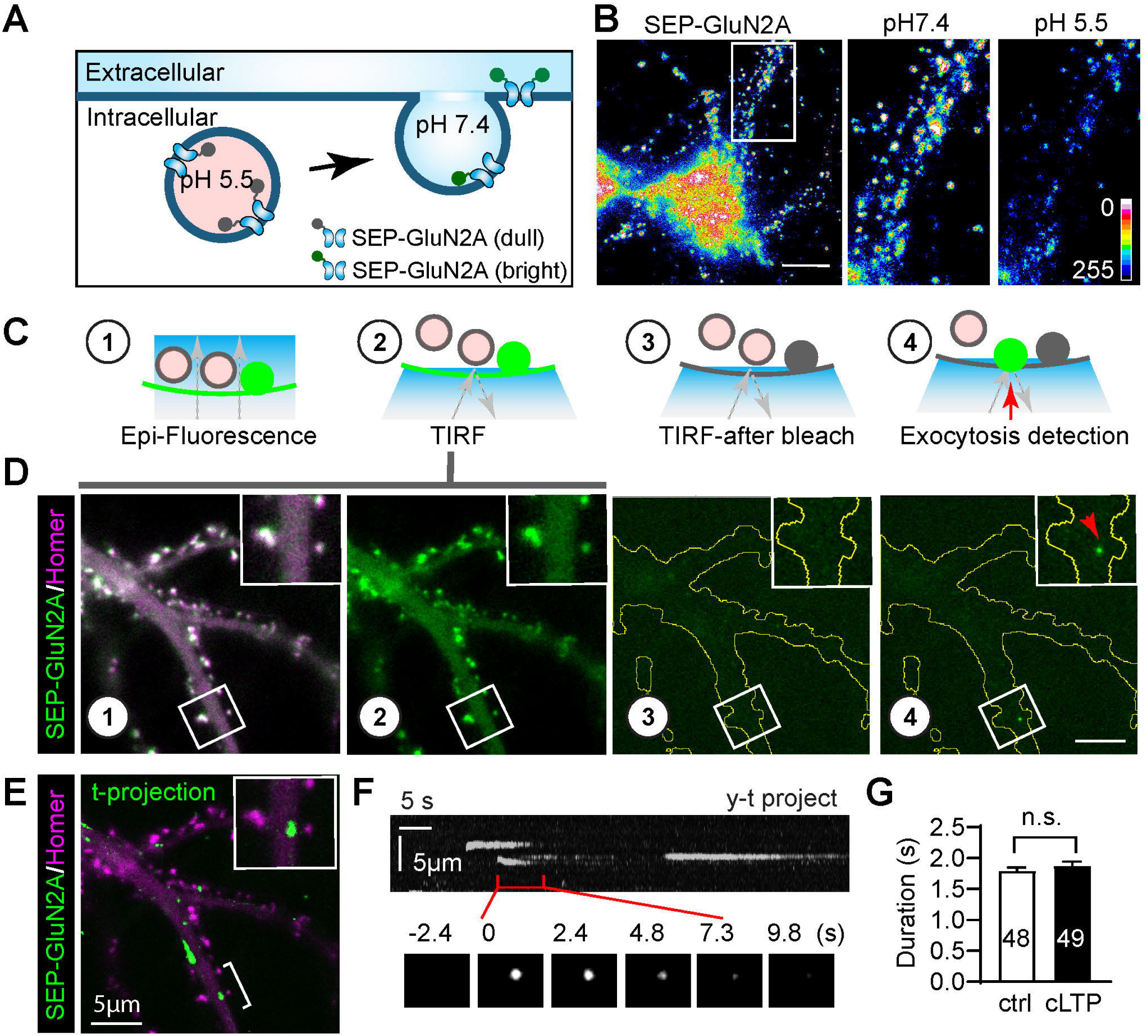
Detection of newly added GluN2A-containing NMDA receptors in dendritic areas of cultured hippocampal neurons. **(A)** Schematic diagram of the mechanism underlying detection of exocytotic pH-sensitive SEP-GluN2 receptors at the neuronal surface. **(B)** Acidification test of SEP-GluN2A subunits expressed in DIV21 hippocampal neurons. pH values of the imaging solution are ∼7.4 and ∼5.5, respectively. Fluorescence intensity is color coded, bar=10 μm. **(C)** Schematic diagram showing bleaching assay in TIRF mode to detect SEP-GluN2 exocytosis. Individual exocytotic events are indicated with red arrow. **(D)** Representative time-lapse images of hippocampal neurons transfected with Homer1c-DsRed and SEP-GluN2A are shown to illustrate the actual detection steps of the bleaching assay for SEP-GluN2 exocytosis (step 1 and 2). Boxed regions are amplified in the upper right corner. The neurons after bleaching are outlined in yellow (step 3). An exocytosis event is annotated with red arrow in step 4. Bar=5 μm. **(E)** Representative maximum projection of the time-lapse image stacks showing exocytic events at 5 min. Bracketed area is amplified in upper right box. Bar=5 μm. **(F)** y-t projection of the region bracketed in **(E)**, with time-lapse frames of an exocytotic event shown in bottom panels. Scale bar as indicated. **(G)** Quantification of average duration of GluN2A exocytic fusion events in neurons before (control) and after 5 to 10 min of cLTP treatment (cLTP). Results are shown in mean±SEM, n.s. no significant difference, n as indicated. Neurons were from three independent preparations, two tailed student’s *t*-test.

**Figure 6.**
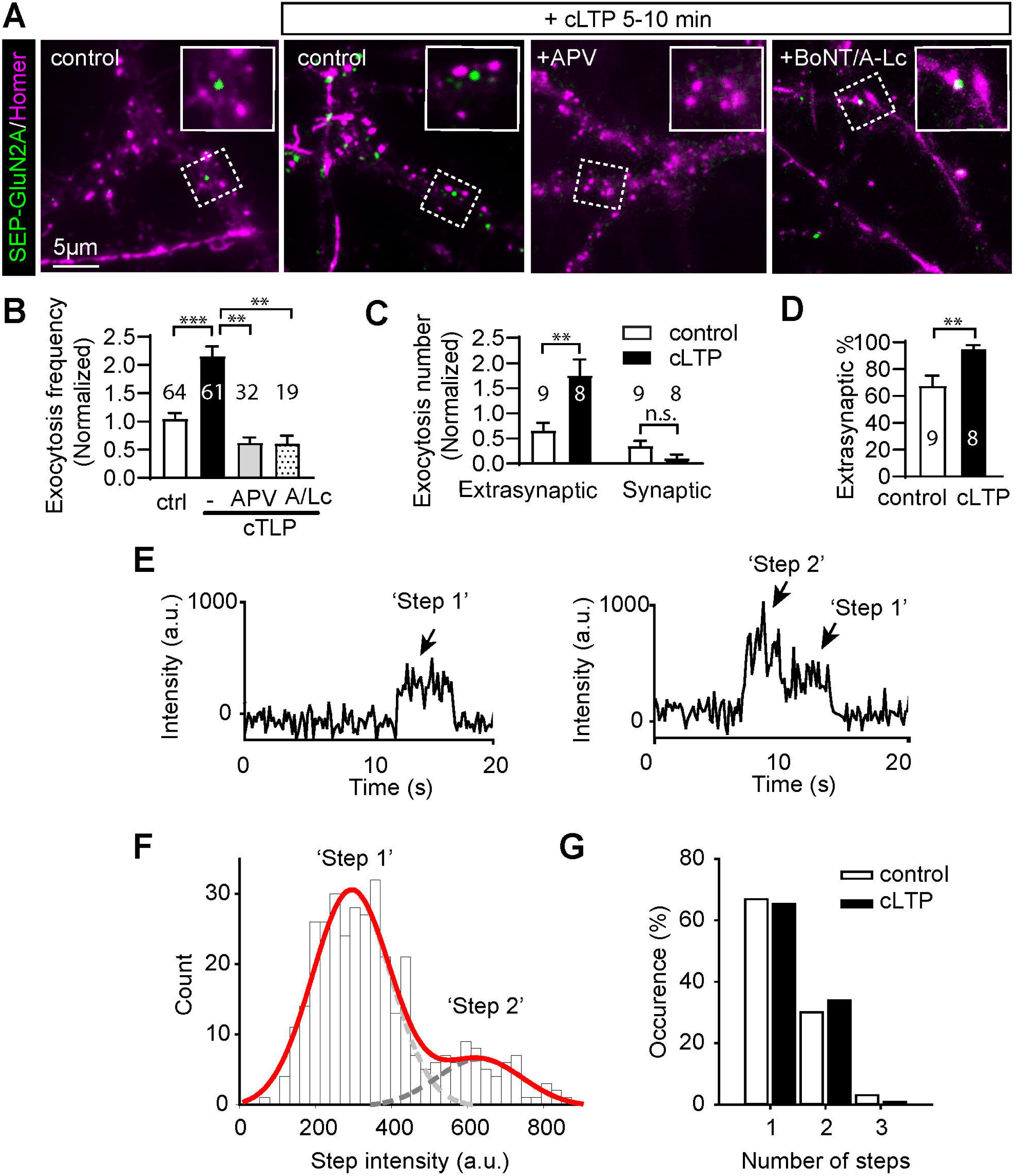
Exocytosis frequency of the GluN2A-containing NMDA receptors in extrasynaptic regions is up-regulated by cLTP. **(A)** Representative figures of cultured hippocampal neurons with indicated treatments showing sites of GluN2A exocytosis. Postsynaptic regions are labelled with Homer-DsRed. Boxed regions showing the exocytic events near the spine are amplified in the upper right corner. Bar=5 μm. **(B)** Quantification of the relative number and **(C)** extrasynaptic/synaptic ratio of SEP-GluN2A exocytosis in neurons before (control) and after 5 to 10 min of cLTP treatment (cLTP). **(D)** Quantification of exocytosis frequency in neurons before (control) and after 5-10 min of cLTP treatment (cLTP), which were compared to those of APV (50 μM)-treated or BoNT/A-Lc (A/Lc) transfected neurons. Results are shown in mean±SEM, \*\**p*<0.01, \*\*\**p*<0.001, n as indicated. Neurons were from at least three independent preparations, two tailed student’s *t*-test. **(E)** Fluorescence intensity recorded from each exocytic fusion event decayed in step-like manner. Each step corresponds to one inserted and bleached SEP-GluN2A subunit. Most intensity-time traces showed one bleaching step (left graph) and smaller number of traces had two bleaching steps (right graph). The steps are indicated with arrows. **(F)** A histogram showing fluorescence intensity distribution of fusion events before step bleaching has occurred. The two peaks are due to traces with one (light gray Gaussian fit) and two (dark gray Gaussian fit) bleaching steps. **(G)** A histogram showing the number of traces that had one, two or three bleaching steps in control (white) and cLTP treated neurons (black). Maximum number of steps observed was three. 163 (control) and 111 (cLTP) exocytic fusion events were analyzed.

As these events were detected after bleaching of pre-existing surface NMDARs, they represent only the newly inserted GluN2A-containing NMDARs occurring upon cLTP perfusion. To further characterize where these cLTP-induced NMDARs were inserted to, we compared the locations of these events with those of co-transfected postsynaptic marker Homer1c-DsRed, which labelled the synaptic areas (Fig. 6A). Using projection of time-lapse images acquired during 5-10 min of cLTP perfusion (Fig. 6A), we found the majority of these newly inserted NMDARs were in the extrasynaptic areas, which were void of Homer1c-DsRed (Fig. 6A, insets). Both the absolute number (Fig. 6C) and the relative percentage (Fig. 6D) of the insertion events in extrasynaptic areas were up-regulated with the cLTP perfusion. The extrasynaptic insertion of GluN2A-containing NMDARs is similar to that of activity-dependent AMPARs, which occurs exclusively in the extrasynaptic region (Yudowski et al., 2007; Lin et al., 2009).

To evaluate whether the increase in insertion frequency is also followed by a change in the number of GluN2A receptors per fusion event, we examined how the fluorescence intensity of each spot changed over time. We observed steps in the intensity-time trace typical for bleaching of single fluorescent proteins (Ulbrich and Isacoff, 2007). Thus, we applied single-step photobleaching, a technique that allowed us to directly count the number of SEP-GluN2A subunits which were inserted into the plasma membrane in each exocytic event. Fig. 6E shows an example of a typical intensity-time trace with one (upper) and two (lower) bleaching steps. We analysed a total of 274 fusion events, 163 of which were in the control neurons and 111 in the neurons in which cLTP was induced. The majority of traces (66%) had only one bleaching step with a fluorescence intensity centred around 303 counts and 32% had two bleaching steps with a total intensity cantered at 630 counts as shown in Fig. 6F. In only 2 % of traces, we counted three steps. When we compared the number of traces with one, two or three bleaching steps in control neurons to those in neurons where cLTP was induced, we did not see any statistical difference (Fig. 6G). These results indicate that the number of SEP-GluN2A subunits inserted per fusion event did not change when cLTP was induced. Since this type of analysis allows us to visualize only labelled GluN2A subunits and not the endogenous ones and because NMDA receptors can contain one or two GluN2A subunits, we were unable to determine the absolute number of receptors that were inserted into the plasma membrane per fusion event. However, we can speculate that there are at least one and at most three NMDA receptors with one labelled GluN2A subunit present in each fusion event and that the number of GluN2A-containing NMDA receptors per endocytic vesicle does not change after cLTP is induced. Furthermore, the data suggests that the cLTP induces a transient increase in extrasynaptic GluN2A-containing receptors which occurs primarily through up-regulation of the SEP-GluN2A receptor insertion frequency into postsynaptic neurons.

## Discussion

### NMDARs are dynamically regulated during synaptic plasticity

The number and subunit composition of NMDA receptor channels at synapses is tightly controlled to ensure synaptic plasticity. Defects in the surface expression of NMDARs disrupt the hippocampal LTP, impairing spatial memory in mice (Sheng and Kim, 2002; Rezvani, 2006), and have also been identified in a range of human cognitive disorders (McQuail et al., 2016; Tannenholz et al., 2016; Avila et al., 2017; Hopf, 2017). Compared to the highly activity-regulated surface AMPARs, NMDARs are regarded as relatively stationary components in the process of synaptic plasticity as the variations in both current amplitude and number of synaptic NMDA channels caused by *in vivo* activation is small (Hestrin, 1992; Petralia et al., 1999) and the absolute number of GluN1 subunits within individual synapses remain relatively constant during the crucial postnatal developmental stages (Petralia et al., 1999). However, accumulating evidence suggests that synaptic NMDARs levels undergo consecutive and regulated synaptic trafficking and retrieval during synaptic remodeling (Lan et al., 2001; Perez-Otano and Ehlers, 2005; Lau and Zukin, 2007; Lau et al., 2010). Moreover, NMDARs are highly dynamic in altering their subunit composition (Bellone and Nicoll, 2007) via regulated lateral movement of existing GluN2A or GluN2B subunits between synaptic and extrasynaptic regions in an activity-dependent manner (Lau and Zukin, 2007; Newpher and Ehlers, 2008; von Engelhardt et al., 2009; Bard and Groc, 2011; Ferreira et al., 2017). Recently, exocytosis of GluN2A and GluN2B receptors was implicated as a possible mechanism underlying GluN2A and GluN2B ratio swap. One study showed that the supply of GluN2A-containing NMDARs to the postsynaptic region is selectively up-regulated by a mechanism dependent on the endoplasmic reticulum chaperon Bip (Zhang et al., 2015). Interestingly, during this process the surface expression of GluN2B-containing NMDARs shows a decline in response to LTP, suggesting that the two subtypes are regulated separately.

We also observed that LTP induces a rapid and different change in the amount of surface GluN2A and GluN2B subunits. We further demonstrated that the up-regulated GluN2A-containing NMDARs inserted in extrasynaptic regions, which are known to be the predominant insertion sites for other ligand-gated ion channels such as AMAPRs (Ulbrich and Isacoff, 2007; Lin et al., 2009) and GABARs (Gu et al., 2016). Similar to these channels, activity-dependent delivery of NMDAR may also be achieved through a two-step mechanism, comprising of: (1) the extrasynaptic receptor exocytosis and (2) the subsequent lateral diffusion of surface receptors between the extrasynaptic and synaptic pools that eventually leads to the balanced proportion of GluN2A and GluN2B in synaptic NMDAR clusters.

### SNARE proteins mediate the rapid extrasynaptic insertion of NMDARs

NMDAR delivery to the plasma membrane can be rapidly enhanced via up-regulation of the SNARE-dependent exocytosis mechanism (Lan et al., 2001). This effect is likely mediated by phosphorylation of proteins associated with NMDARs (Lan et al., 2001). SNAP-25 and its postsynaptically enriched homologue SNAP-23 are such candidate proteins. Both can be phosphorylated in an activity-dependent manner (Hepp et al., 2005; Lau et al., 2010), and it is likely that the rapid LTP-triggered insertion of NMDARs onto the neuronal surface, as observed in acute brain slices (Grosshans et al., 2002), is driven by SNAP-25 or SNAP-23. Indeed, SNAP-25 was found to be required for the activity-dependent insertion of both NMDARs and AMPARs (Jurado et al., 2013). SNAP-23 is required for the consecutive (basal) surface delivery of NMDARs in mice brain (Suh et al., 2010). However, one recent study found that the insertion of NMDARs, as reflected by the exocytosis of GluN1 subunits, is not regulated by synaptic activation/inhibition paradigms (Gu and Huganir, 2016). Therefore, whether SNAP-23 and SNAP25 directly control the exocytosis of NMDARs during the synaptic modification process is still arguable.

This debate is partially due to the fact that, in most studies, exocytosis of NMDARs was estimated using the surface level of GluN1 subunits, which is the relatively stationary component of NMDARs (Perez-Otano and Ehlers, 2005; Newpher and Ehlers, 2008). Other technical difficulties to overcome include the small number of NMDARs within each exocytosis package, which limits their precise detection in live neurons. We circumvented these limitations by using the TIRF-based bleaching assay to remove most of the preexisting surface GluN2A-containing NMDARs, therefore boosting the detection of newly inserted ones. We further paired this detection with diverse perfusion conditions to examine the impact of synaptic plasticity on this exocytosis process. By combining the bleaching assay with live-cell surface staining, we quantified the cLTP-induced vesicle insertion of GluN2A-containing NMDARs and noticed that the exocytosis frequency of GluN2A-containing postsynaptic vesicles was significantly increased following LTP. We therefore found that postsynaptic exocytosis machinery directly regulates the surface expression of NMDARs in an activity-dependent manner. This selective insertion of GluN2A-containing NMDARs together with their lateral diffusion that happens on longer timescales, controls the number and subunit composition of synaptic NMDARs and thereby orchestrates the synaptic plasticity underlying learning and memory formation.

In conclusion, our study provides direct evidence that NMDAR-mediated LTP induces a transient up-regulation of SEP-GluN2A-containing NMDARs in extrasynaptic sites of postsynaptic neurons. SNAP-23, and also to a minor extent SNAP-25, are required to control regulated surface insertion of GluN2A receptors by controlling their exocytotic frequency at extrasynaptic regions. Either SNAP-23 or SNAP-25 also mediates the consecutive GluN2A membrane delivery. By directly visualizing cLTP-induced SEP-GluN2A exocytotic fusion events, we found that cLTP specifically raised the frequency of GluN2A exocytosis into the extrasynaptic area, without affecting duration or package size of individual insertion events. Our work reveals a novel mechanism underlying the initial NMDAR-mediated postsynaptic response, providing potential methods to specifically abolish regulated NMDA receptor trafficking by interfering with its fusion machinery during LTP.

## Acknowledgments

This work was supported by grants from the Australian Research Council (ARC) DE170100546 to TW; XY were recipients of University of Queensland Research Training Scholarships. We would like to thank Nela Durisic for consistent support and help with this project. We thank the Advanced Microscopy Facility and Animal Facility of QBI for technical support. We are grateful to Victor Anggono, Frederic Meunier for generously providing their lab resources for this study. We would like to thank Men chee Tan, Se-Eun (Joanne) Jang, He Huang, Christopher Small, Dejan Gagoski and Xuanling Hillary Yong for their work in preparing plasmids and neurons for this study. We are grateful to Brett Collins for his constructive discussion in the initial stage of the study. We thank Russell Wilson for proof reading and Jane Mooney for critical reading of the manuscript.

---

**Supplementary Figure 1.**
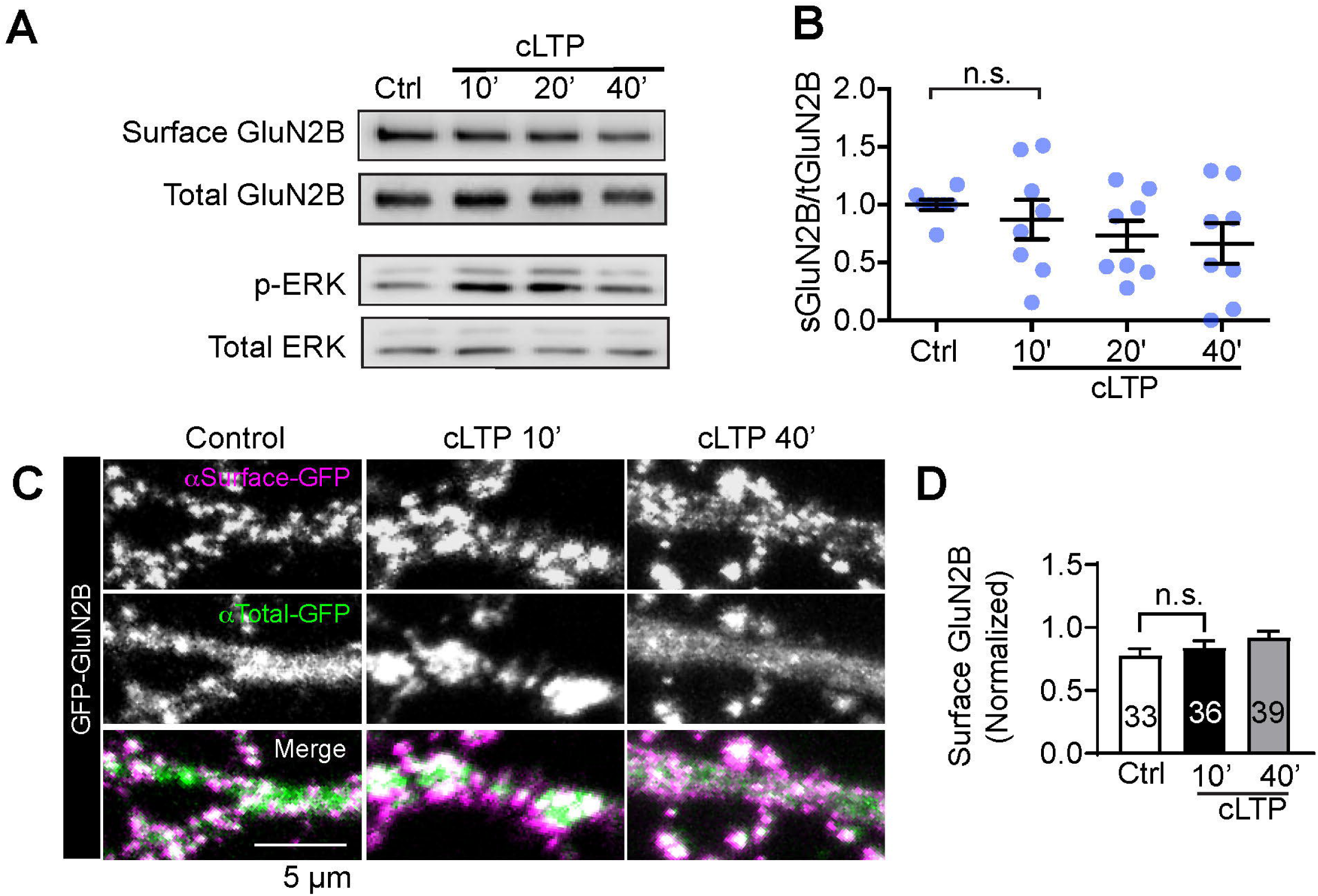
cLTP induces a transient increase in surface GluN2A (but not GluN2B) subunits of NMDA receptors. **(A)** Representative western blots showing the levels of biotin labeled surface and total GluN2B receptors in cultured rat cortical neurons, treated as indicated. The relative level of phosphorylated ERK (pERK) is used as the marker for cLTP induction. **(B)** Quantification of **(A)**. Results are shown in scatter plot with mean±SEM, n.s. *p*>0.05. n=8 repeats from eight independent preparations, two tailed student’s *t*-test. **(C)** Representative surface immunostaining of extracellular GFP tag in GFP-GluN2B expressing hippocampal neurons treated as indicated. Boxed regions are amplified in lower panels. Bar=5 μm. **(D)** Quantification of **(C)**. Results are shown in mean±SEM, n.s. *p*>0.05. N represents the number of cells analyzed and are shown on the bar. Data were from three independent cultures, two tailed student’s *t*-test.

**Supplementary Figure 2.**
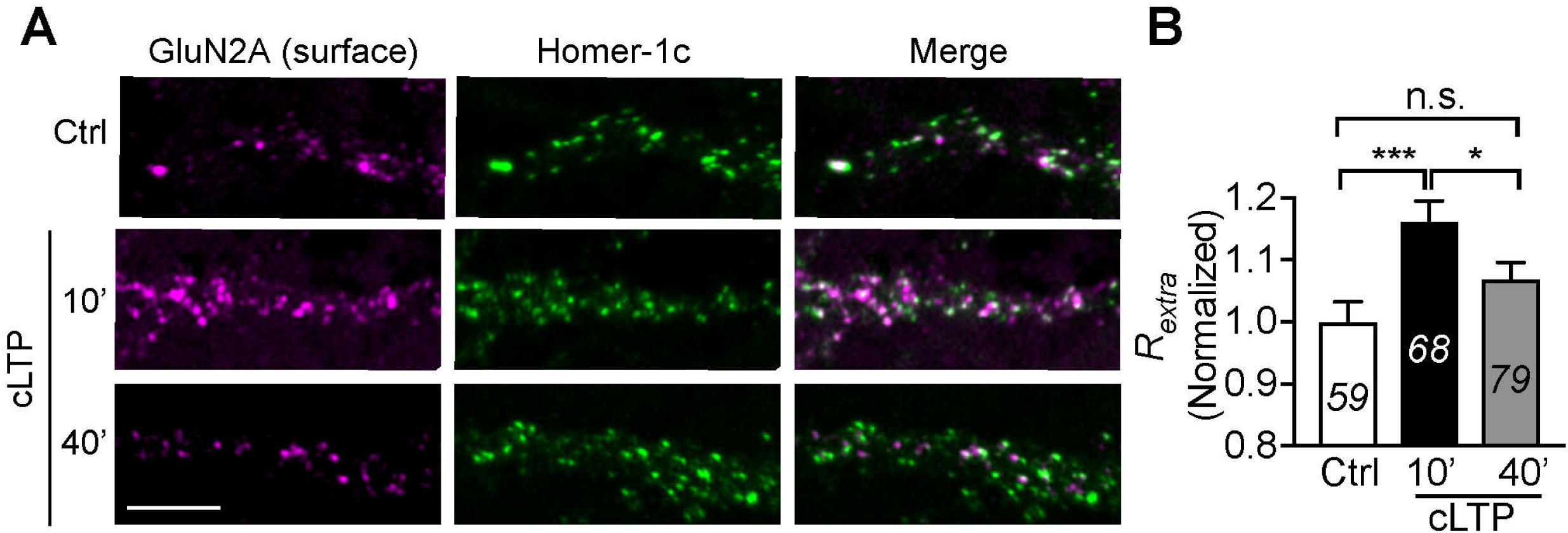
cLTP up-regulates the ratio of surface GluN2A receptors in the extrasynaptic region. **(A)** In DIV21 rat hippocampal neurons, surface GluN2A NMDA receptors were stained with specific antibodies against their extracellular domain before fixation, followed by immunostaining using Homer-1c antibody. Confocal microscopy was used to visualize the distribution of fluorescent signals at dendritic spines. Scale bar= 5 μm. **(B)** The ratio of GluN2A receptors localized in the extrasynaptic region was defined as the *R_extra_*, which was calculated from the ratio of GluN2A not overlapping with Homer-1c (See Methods). Data represent mean±SEM, \**p*<0.05, \*\**p*<0.01. N represents the number of cells analyzed and are shown on the bar. Data were measured from two independent cultures, two-tailed student’s *t*-test.

**Supplementary Figure 3.**
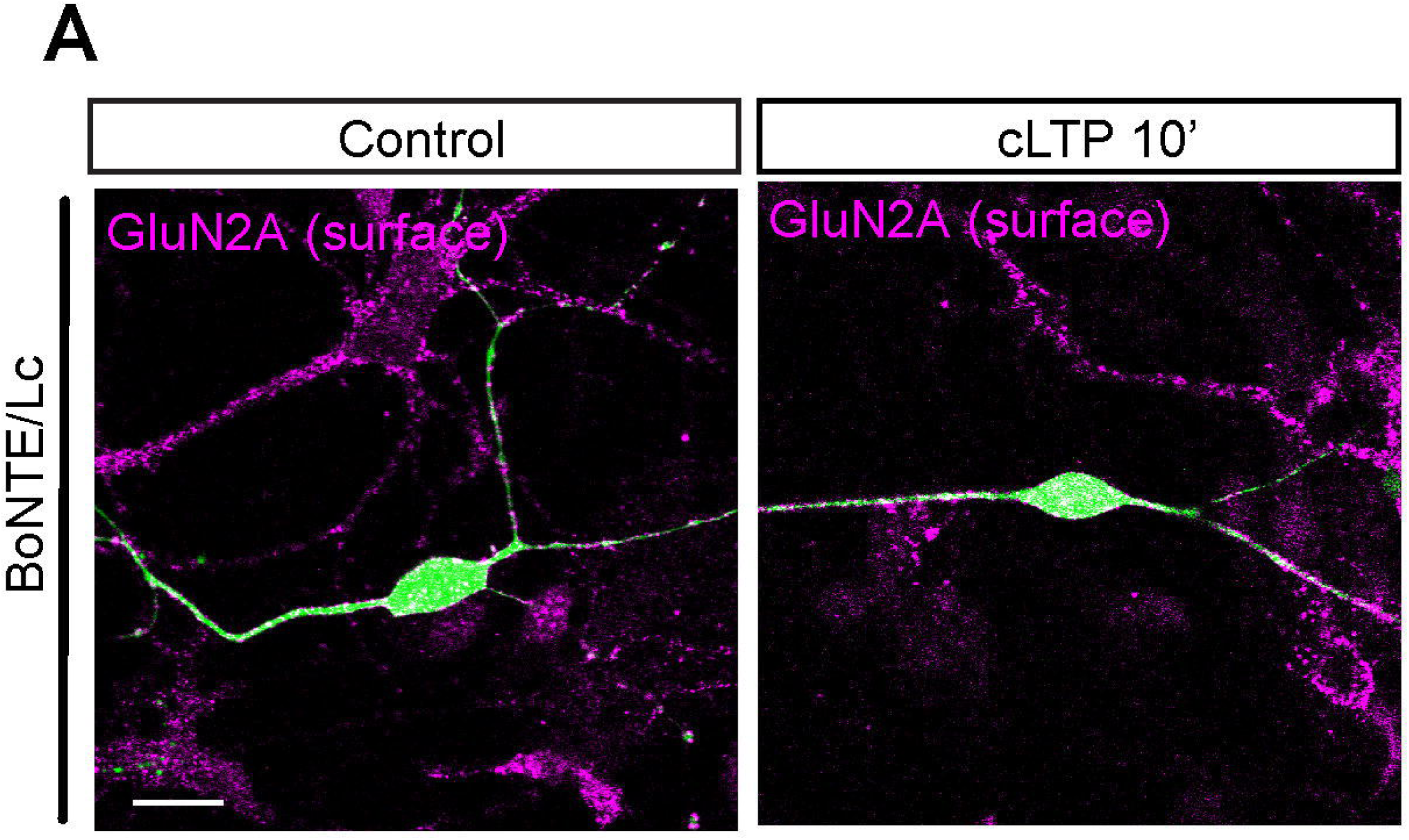
Neuronal degeneration caused by the expression of BoNT/E-Lc in postsynaptic neurons. Mature rat hippocampal neurons were transfected with BoNT/E-Lc-GFP on DIV14-17, treated with cLTP stimulation on DIV21, followed by surface GluN2A immunostaining before fixation. Representative confocal images show the typical neurodegeneration phenotype of beading axons and round cell bodies in all BoNT/E-Lc-GFP expressing neurons treated as indicated. Bar = 20 μm.

**Supplementary Figure 4.**
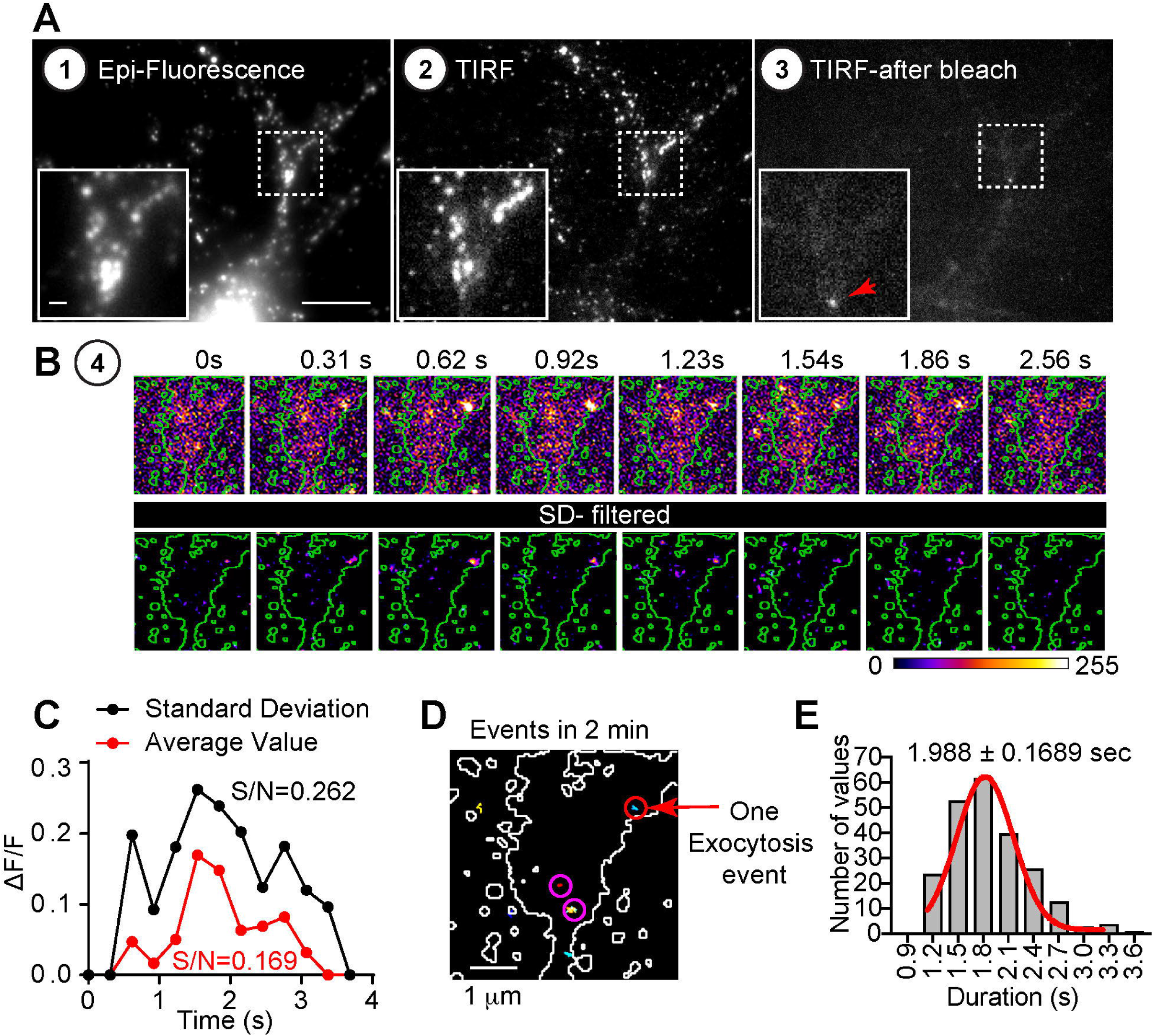
Automatic detection of SEP-GluN2A exocytosis events in the dendritic regions of cultured hippocampal neurons. **(A)** Representative images showing each step of the bleaching assay for the automatic analysis of exocytosis. Boxed regions are amplified in the lower left corner. Bar= 10 μm. Inset Bar = 1 μm. **(B)** The last step (Step 4) of the image preparation of the boxed region in **(A)** shows effects of standard deviation filter (see Methods). For comparison, time-lapse images of unfiltered (top panels) and SD-filtered (bottom panels) are shown. **(C)** Comparison of improvement in signal to noise ratio averaging of image sequence (red) or when SD-filter was applied (black). **(D)** The trajectories detected using TrackMate plugin of Image J. One exocytosis event is marked out with red arrow. Bar as indicated on the figures. **(E)** Histogram of the average duration of SEP-GluN2A exocytotic trajectories detected by automatic method. The average duration of control condition is shown as mean ± SEM, from n=50 neurons of three different preparations. R^2^=0.983 with the least squares Gaussian curve fitting method.

**Supplementary Movie 1. cLTP treatment increases the intensity of Calcium current in the dendritic spines of cultured hippocampal neurons.**

Movies showing the calcium signal fluctuation in the dendritic spines of of dendritic spine regions visualized in GCaMP6-expressing hippocampal neurons, which were treated before (control) or during the cLTP (+cLTP). Fluorescence intensity of boxed area is amplified with fluorecent intensity colour-coded in lower boxes. Bar=20 μm.

**Supplementary Movie 2. GluN2A containing NMDAR Exocytosis detected using SEP-GluN2A transfected neurons.**

Representative movie of hippocampal neuron transfected with Homer1c-DsRed and SEP-GluN2A are shown to illustrate the actual SEP-GluN2 exocytosis. Boxed regions are amplified in the lower pannels. The neurons were imaged following a bleaching step are overlayed with the Homer-DsRed signal in the right pannels. Bar=5 μm.

## Author contributions

X.Y. prepared primary neuronal cultures, conducted most of the biotinylation and western experiments. T.W. performed TIRF and SIM microscopy assays, analyzed the results and drafted the manuscript. W.L. performed and analyzed live-imaging and surface staining data.

## Materials & correspondence

Correspondence and requests for materials should be addressed to Dr Tong Wang.

## Data Availability

The data that support the findings of this study are available on request from the corresponding author Tong Wang.

